# The CCR4/CCL17 axis drives intestinal acute Graft versus Host disease after allogeneic bone marrow transplantation

**DOI:** 10.1101/2024.03.02.583093

**Authors:** Sebastian Schlaweck, Michelle Klesse, Oliver Schanz, Selina K. Jorch, Glen Kristiansen, Marieta Toma, Chrystel Flores, Janine Becker-Gotot, Christian Kurts, Peter Brossart, Dominik Wolf, Annkristin Heine

**Affiliations:** Medical Clinic III for Hematology, Oncology, Rheumatology, Immunoncology and Stem-cell transplantation, University of Bonn; Institute for Pathology, University of Bonn; Institute of Molecular Medicine and Experimental Immunology, University of Bonn; Internal Medicine V, Department of Hematology and Oncology, Comprehensive Cancer Center Innsbruck (CCCI), Tyrolean Cancer Research Center (TKFI), Medical University Innsbruck (MUI), Innsbruck, Austria; Mildred Scheel School of Oncology Aachen Bonn Cologne Düsseldorf (MSSO ABCD), Bonn, Faculty of Medicine and University Hospital of Bonn, D-53127, Bonn, Germany

**Author notes:** **Correspondence:** Annkristin Heine, MD, Medical Clinic III, University Hospital Bonn, Venusberg Campus 1, 53127 Bonn, Germany.

**Keywords:** Acute Graft versus Host disease, CCR4, CCL17, TH2, Gata3, ruxolitinib

## Abstract

Acute-Graft-*versus*-Host disease (aGvHD) is a life-threatening complication after allogeneic stem-cell-transplantation. It is mediated by alloreactive T cells whose trafficking to aGvHD target organs is orchestrated by chemokines.

We here asked whether CCL17 and its corresponding receptor CCR4 are involved in aGvHD development and severity. We applied an experimental mouse model of aGvHD in CCR4/CCL17 knockout mice and analyzed gut biopsies of GvHD patients.

We show that the absence of CCR4 in transplanted T cells induced significantly less severe aGvHD. This was accompanied by reduced expression of Gata3. Mechanistically, only CD4^+^, but not CD8^+^CCR4^-/-^ T cells protected from aGvHD. We next identified dendritic cells in the small intestine to produce CCL17, which selectively recruited CD4^+^ T cells. IL-4 production by intestinal CD4^+^ T cells promoted proliferation of CD8^+^ T cells. In line, we detected an upregulation of CCL17 and *Gata3* in human aGvHD samples.

Our results indicate that local CCL17 production in aGvHD target organs recruits T cells, reinforcing local tissue damage and immune cell recruitment. We identified the JAK1/2-inhibitor ruxolitinib to dampen CCL17-expression, thereby reducing GvHD severity.

We here dissect a to date unknown role of the CCL17-CCR4 axis in aGvHD, which might help to develop novel therapeutic strategies.

## Introduction

Allogeneic hematopoietic stem cell transplantation (alloHSCT) is the only curative treatment for many hematological malignancies. However, serious life-threatening complications, such as acute Graft-versus-Host disease (aGvHD), limit its success. AGvHD is mediated by allo-reactive donor T cells and occurs in up to 50% of patients undergoing alloHSCT, with approximately 15% of patients developing severe steroid-refractory aGvHD ^1^. The latter is currently associated with high mortality rates and limited available treatment options. The recent approval of the JAK1/2 inhibitor ruxolitinib for both steroid-refractory acute and chronic GvHD has expanded the armamentarium for this difficult-to-treat patient population ^2,3^, the demand for additional preventative or therapeutic strategies remains high.

Prior to transplantation, patients receive cytotoxic conditioning therapy to dampen their immune system and facilitate stem cell engraftment. However, conditioning also induces tissue damage in healthy organs and barrier sites, resulting in the release of damage-associated molecular patterns (DAMPs), such as adenosine-tri-phosphate (ATP), interleukin-33 (IL-33), uric acid, and high mobility group box 1 protein (HMBG-1). Innate immune cells are among the first to respond to conditioning-induced injury^4^. They migrate into secondary lymphoid organs upon activation and facilitate the priming of allogeneic donor T cells, which then infiltrate GvHD target organs (skin, liver, and GI tract) and promote further tissue damage^5^.

Chemokines are essential for orchestrating immune cell trafficking under homeostatic and pathological conditions. In doing so, they are involved in developmental and maturation processes; wound healing; and the initiation, maintenance, and regulation of robust immune responses^6,7^. Several pro-inflammatory chemokines and their receptors have been implicated in aGvHD, which is characterized by extensive immune cell migration ^8^. For example, CCL5 and CXCL10 have been shown to mediate T-cell infiltration into lymphoid tissues and target organs^9,10^. Conversely, blockade of the chemokine receptor CCR5 confers protection against GvHD in patients following reduced-intensity alloHSCT ^11^. These data suggest a promising therapeutic potential for targeting chemokines and their receptors in aGvHD.

The CCL17-CCR4 axis has not been explored in aGvHD. Previous studies have demonstrated that CCL17 is relevant to various autoimmune diseases and is associated with the pathogenesis of intestinal inflammation and murine sclerodermatous chronic GvHD ^12–15^. CCR4 is a high-affinity receptor for CCL17 and CCL22, and is predominantly expressed on T helper (TH) 2, skin-homing T cells, and regulatory T cells (Tregs) ^16,17^. CCR4^+^ T cells contribute to tissue damage after solid organ transplantation and in diabetes ^18,19^.

Therefore, we propose that CCL17/CCR4-guided T cell migration may also play a role in GvHD, particularly in intestinal aGvHD. Here, we provide evidence that the expression of CCR4 by allogeneic donor T cells and CCL17 in the recipient indeed correlates with aGvHD severity in mice. Our data suggest that the CCL17-CCR4 axis may represent a novel therapeutic strategy for the prevention and treatment of aGvHD and could also aid in optimizing treatment for steroid-refractory GvHD.

## Methods

### Human subjects

This study was conducted in accordance with the Declaration of Helsinki, and approval was obtained from the Institutional Ethics Committee of the University of Bonn (#175/20). Endoscopic and histological examinations of patients undergoing alloHSCT at the University Hospital of Bonn were performed according to clinical routine. CCL17 antibody (goat anti-CCL17, Sigma Aldrich, #C1497) and Gata3 antibody (Biocare, #L50-823) were used for immunohistochemistry. Representative parts of the sections were selected for immunohistochemical analysis in a semi-quantitative manner using ImageJ Version 2.1.0/1.53c (http://imagej.net). Patient characteristics are listed in Supplementary Table 1.

### Mice and transplantation model

All animal experiments were approved by the *Landesamt für Natur, Umwelt und Verbraucherschutz Nordrhein-Westfalen (LANUV)*. Six-to twelve-week-old Balb/c mice received total body irradiation (9 Gy, 2 split doses, time between each dose was at least 4 h) the day before transplantation. Bone marrow (BM) cells from C57BL/6 donor mice (matching age and sex) were T cell-depleted using CD90.2 beads (Miltenyi, #130-121-278). Splenic T cells were isolated using the CD3e MACS (Miltenyi, #130-094-973) from either wildtype (wt) or CCR4^-/-^ animals (C57BL6/ background, matched in age and sex). T cell purity was always >90%. Balb/c mice received either 5 × 10^6^ BM cells alone (no GvHD control) or together with 6 × 10^5^ T cells for aGvHD induction. In mixed T-cell transplantation experiments, CCR4^-/-^ and wt CD4^+^ and CD8^+^ T cells were mixed in a physiological 3:2 ratio.

When indicated, ruxolitinib treatment was administered as previously described^20^. Briefly, mice were fed either ruxolitinib (Selleckchem, #S1378, 75 mg/kg, dissolved in 0.1% Carboxy-Methyl-Cellulose (CMC)) or vehicle twice daily beginning on the day of irradiation. Mice were sacrificed on day 3 after transplantation. Organs were harvested and the small intestines were analyzed for chemokine expression by real-time quantitative PCR(qPCR).

The contribution of CCL17 was investigated using either C57BL/6 (age 8-12 weeks) or CCL17^-/-^ recipient mice (on a C57BL/6 background). The mice received total-body irradiation 24 h before transplantation. BM and T cells were isolated as previously described. Recipient mice received 5 × 10^6^ BM cells and 1 × 10^6^ T cells from Balb/c donor mice.

### Isolation of dendritic cells and *in vitro* stimulation

Splenic immune cells were isolated from explanted organs of C57BL/6 mice. The organs were minced, and single-cell suspensions were washed. Dendritic cells (DC) were isolated using CD11c microbeads (Miltenyi, #130-125-835), according to the manufactureŕs instructions. DC were cultured in RPMI 1640 medium supplemented with 10% FBS, 1% P/S for 24h in the presence of IL-33 (Peprotech, #210-33, final concentration 100ng/ml). When indicated, ruxolitinib (Selleckchem, #S1378) was added before IL-33 stimulation at a final concentration of 1µM or 10µM. Chemokine secretion was investigated by analyzing the supernatants with the LEGENDplex Mouse Proinflammatory Chemokine Panel (BioLegend #740007) according to the manufacturer’s instructions.

### Quantitative reverse transcriptase polymerase chain reaction

Mice were sacrificed at the indicated time points and tissues were snap-frozen in liquid nitrogen. Alternatively, cells were harvested from *in vitro* culture. RNA was extracted using RNAzol (Sigma Aldrich, #R4533) and cDNA was synthesized using the RevertAid First Strand cDNA Synthesis Kit (Thermofisher, #18091050) according to the manufacturer’s instructions. Quantitative real-time PCR (qPCR) was performed using specific primers (0.2µM concentration and SYBR Green PCR Master Mix (Applied Biosystems, #4309155) on the Mastercycler RealPlex2 (Eppendorf). Data were normalized to GAPDH expression and analyzed using the ΔΔCT method. Primer sequences are listed in Supplementary Table 2.

### Murine CCL17 expression

For immunohistochemistry, tissues were embedded in Tissue Tek O.C.T. compound (Plano, #R1180-X) and cut into 5 µm slices. The slides were stained with antibodies against CD11c (clone N418) and DAPI.

To assess the expression of CCL17, areas were drawn and measured along the whole tissue comprising the mucosa, including villi and crypts, excluding the submucosa and muscularis propria. CCL17 expression areas within the whole tissue were measured and normalized to the whole tissue in mm²). Images were acquired using a ZEISS LSM710 Observer and corresponding ZEN Black software.

### Cytokine measurement

Blood was collected on day 7 after transplantation *via* a puncture of the tail vein. Serum was obtained and the Th1/Th2 Cytokine 11-Plex Mouse ProcartaPlex™ Panel (Invitrogen, EPX110-20820-901) was used according to the manufacturer’s instructions to quantify cytokine levels.

### Histological examination

Seven days after alloHSCT, sections of the small intestine, large intestine, and liver were collected, fixed in 4% paraformaldehyde (PFA) and stained with hematoxylin/eosin (HE). Acute GvHD was scored blinded to the treatment groups according to a previously published histopathology scoring system^33^.

### Flow cytometry

T cells were isolated from the gut as previously described ^21^. Briefly, the small intestine was removed from feces and washed in a mixture containing HBBS + 2%FCS + 0.5 M Dithiothreitol (DTT). Tissue was digested with 2mg/ml Collagenase in RPMI+ 2% FCS for one hour at 37°C in an incubator (5% CO2). The homogenized sample was rinsed through a 70 µm mesh and the cell suspension was stained with the respective antibodies listed in supplementray Table 2. For cytokine measurement, cells were re-stimulated as described below and measured with a BD FACSCanto II flow cytometer. Subsequently, data were analyzed using FlowJo Software v10.7.1 (Treestar, Ashland, USA).

### Transwell assay

Splenic T cells were isolated by CD3e MACS kit (Miltenyi #130-094-973) from mice injected with 0.2 µg α-GalCer (1 nmol; Axxora, #ALX-306-027) 24h before. 600 µl of RPMI 1640 medium supplemented with 10% FBS, 1% P/S containing CCL17 (800ng/ml, R&D Systems, #529-TR), when indicated, was added to a well of a 24 well plate and a 6.5 mm Transwell® Polycarbonate Membrane insert with a 5.0 µm pore (Costar #3421/Sigma # CLS3421-48EA) was added to each well. One hundred microliters of T-cell suspension (1 × 10^6^ /ml) was carefully added to the insert without touching the membrane. cells were incubated for six hours at 37°C and 5% CO2. After carefully removing the inserts, the migrated cells were collected from each well, quantified, and characterized by flow cytometry. Therefore, cells collected from the lower chamber were stained with FACS antibodies. Cells were counted for 90 s using a BD FACSCanto II flow cytometer.

### T cell differentiation and intracellular staining

Naïve CD4^+^ or CD8^+^ T cells were isolated using a MACS system and the respective isolation kits (Miltenyi, #130-104-453 and # 130-096-543) according to the manufacturer’s protocol. For T cell proliferation, cells were seeded in RPMI 1640 medium supplemented with 10% FBS, 1% P/S, and 0.1% β-mercaptoethanol in the presence of αCD3/CD28 beads (Thermofisher, #11456D). The stimulants for Th differentiation, stimulation, and staining are listed in Supplementray Table 2. For Th polarization, cells were seeded in supplemented RPMI 1640 medium, and cytokines were added as described in Supplementary Table 2. When indicated, CCL17 (800ng/ml, R&D Systems, #529-TR) was added during the differentiation process. For analysis, cells were re-stimulated (Supplementary Table 2) for 4 h and stained for intracellular expression of either IL-4 or FOXP3 after fixation and permeabilization using the Fixation Buffer (BioLegend, # 420801), the Intracellular Staining Perm Wash Buffer (BioLegend, # 421002), or the Foxp3/Transcription Factor Staining Buffer Set (eBioscience, #00-5523-00).

### Statistical analysis

Data were analyzed using GraphPad Prism Version 9.4.0. Grubb’s test with α=0.05 was performed to identify outliers. The Student’s t-test was performed to compareing two groups. When comparing more groups, an ordinary one-way ANOVA with Šídák’s multiple comparison test was used. Error bars represent standard deviation. p value below 0.05 was considered depicted significant and is depicted as *. P values <0.01 are depicted as ** and <0.001 as ***. Survival experiments were analyzed using the Mantel-Cox test.

## Results

### Murine GvHD depends on CCR4 expression on allogeneic donor T cells

Chemokine receptors orchestrate T-cell migration in aGvHD. Studies elucidating the role of CCR5 and CXCR1^10,11^, for example, have led to the development of therapeutic approaches to prevent aGvHD^22^, but data on the relevance of CCR4 in aGvHD are scarce. Previous studies highlighting the importance of CCR4 and CCL17 in intestinal inflammation^13^ and alloreactivity^19^ have suggested that the CCR4/CCL17 axis could also be a promising target for the treatment of aGvHD.

In this study, we employed an established mouse model^23^ for experimental aGvHD, which is based on an MHC mismatch (C57BL/6 → BALB/c). To elucidate the role of CCR4 in GvHD initiation, we transplanted recipients with wt BM and either CCR4^+/+^ (wt) or CCR4^-/-^ T cells (Fig. 1A, left). While wt mice showed typical signs of GvHD, such as loss of weight, diarrhea, and loss of fur, we observed prolonged survival in mice that received CCR4^-/-^ T cells compared withto wt T cells (Fig. 1A, p=0.0158 and Supplementary figure 1A). In line with these findings, histopathological GvHD scores of the liver and the small and large intestines were also reduced (Fig1. B + C, large intestine (LI, p=0.0484), small intestine (SI, p=0.0174), and liver (LIV, p=0.0126)). In addition, recipients of CCR4^-/-^ T cells displayed decreased serum levels of pro-inflammatory cytokines such as IL-13, IL-12p70, INFγ, and GM-CSF (Fig. 1D). These data show for the first time that CCR4 deficiency in donor T-cells reduces the risk of GvHD.

**Figure 1.**
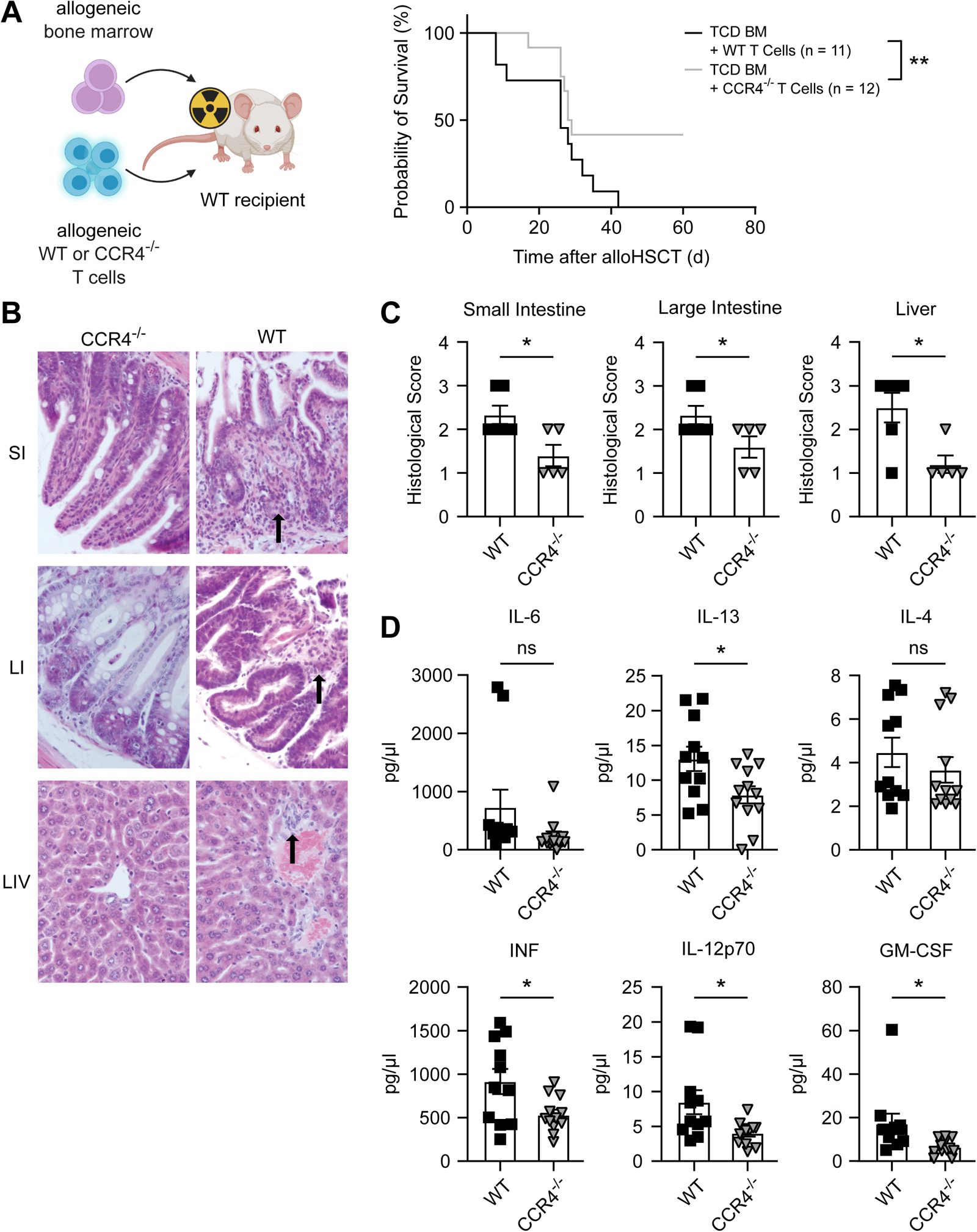
Transplantation of CCR4-/-T cells ameliorates experimental aGvHD. (A) Balb/c mice were irradiated and received bone marrow from C57BL6/N donor animals. Survival was analyzed and recipients of CCR4-/-T cells, in comparison to wt T cells, showed significantly prolonged survival (*p=0.0158)*. Schematic figure created using biorender.com. (B) Representative sections of small intestine (SI), large intestine (LI) and liver (LIV) were isolated on day 7 after alloHSCT from mice transplanted with CCR4-/-or WT T cells. (C) Histopathologic changes (large intestine (*p=0.0484*), small intestine (*p=0.0174*) and liver (*p=0.0126*)) show GvHD specific pathology in recipients of *wt* T cells. (D) Serum cytokine levels were measured and compared between recipients of *wt* and CCR4-/-T cells. Cytokine levels for IL-6 (*p=0.1075*), IL-13 (*p=0.0241*), IL-4 (*p=0.3794*), IL-12p70 (*p=0.0178*), INFγ (*p=0.0193*) and GM-CSF (*p=0.0247*) were altered between both groups.

### CCR4^+^ CD4^+^ T cells drive intestinal aGvHD

CCR4 is expressed by T helper 2 (TH2) cells^24^. We analyzed the RNA expression levels of signature transcription factors for Th subsets in the terminal ileum of mice that received either wt or CCR4^-/-^ T cells and found that only *Gata3*, the key transcription factor for TH2 cells, was differentially expressed (Fig. 2A, expression of *Tbet*, *FOXP3* and *Rorc* (not significant) and *Gata3* (p=0.0491)). *Gata3* expression was significantly higher in recipients of wt than in CCR4^-/-^ T-cells. The data were supported by increased IL-4 production in intestinal CD4^+^ T cells from recipients of T cells compared to CCR4^-/-^ T cells (Fig. 2B, p=0.0031).

We subsequently analyzed gut biopsies from patients with suspected intestinal aGvHD and late-onset aGvHD after alloHSCT for the expression of *GATA3* (Fig. 2C). The median time point at which biopsy was performed in patients after alloHSCT was 95.46 days [range, 21–222 days] in the GvHD group and 114.55 days [range, 20–217 days] in the non-GvHD group (patient characteristics are listed in Supplementary Table 1). Relative frequencies of *GATA3^+^* cells were significantly higher in patients with aGvHD grades II-IV compared than in those without GvHD (Fig. 2D, p=0.0028), thus corroborating our *in vivo* experiments.

Based on these results, we performed transplantation experiments with a mixed CCR4^+/+^/^-/-^ T cell compartment (CD4^+^ CCR4^-/-^/CD8^+^ wt or CD4^+^ wt/CD8^+^ CCR4^-/-^). Notably, survival was only improved when mice received CCR4^-/-^ CD4^+^ T cells, either with CCR4^-/-^ or wt CD8^+^, but not when CCR4 was solely lacking in the CD8^+^ compartment (Fig. 2E, p= 0.0009 and Supplementary figure 1B for weight curves).

### CD8^+^ T cells do not express CCR4 but their proliferation is stimulated by T_H_2 cytokines

To further decipher how CCR4^+^CD4^+^ T cells contribute to aGvHD development, we examined T cell composition in the small intestine on day 7 after transplantation and observed higher frequencies of CD4^+^ T cells in recipients of CCR4^-/-^ compared to wt T cells (Fig 3A, p= 0.0038). Consequently, frequencies of CD8^+^ T cells were significantly lower (Fig. 3B, p=0.005), coinciding with a decrease in the relative mean fluorescence intensity of Ki67 in CD8^+^ T cells, indicating diminished proliferation rates (Fig. 3B, p=0.0097). In contrast, the relative frequencies of Tregs, NKT cells, and NK cells did not change (Supplementary Fig. 1C).

**Figure 2.**
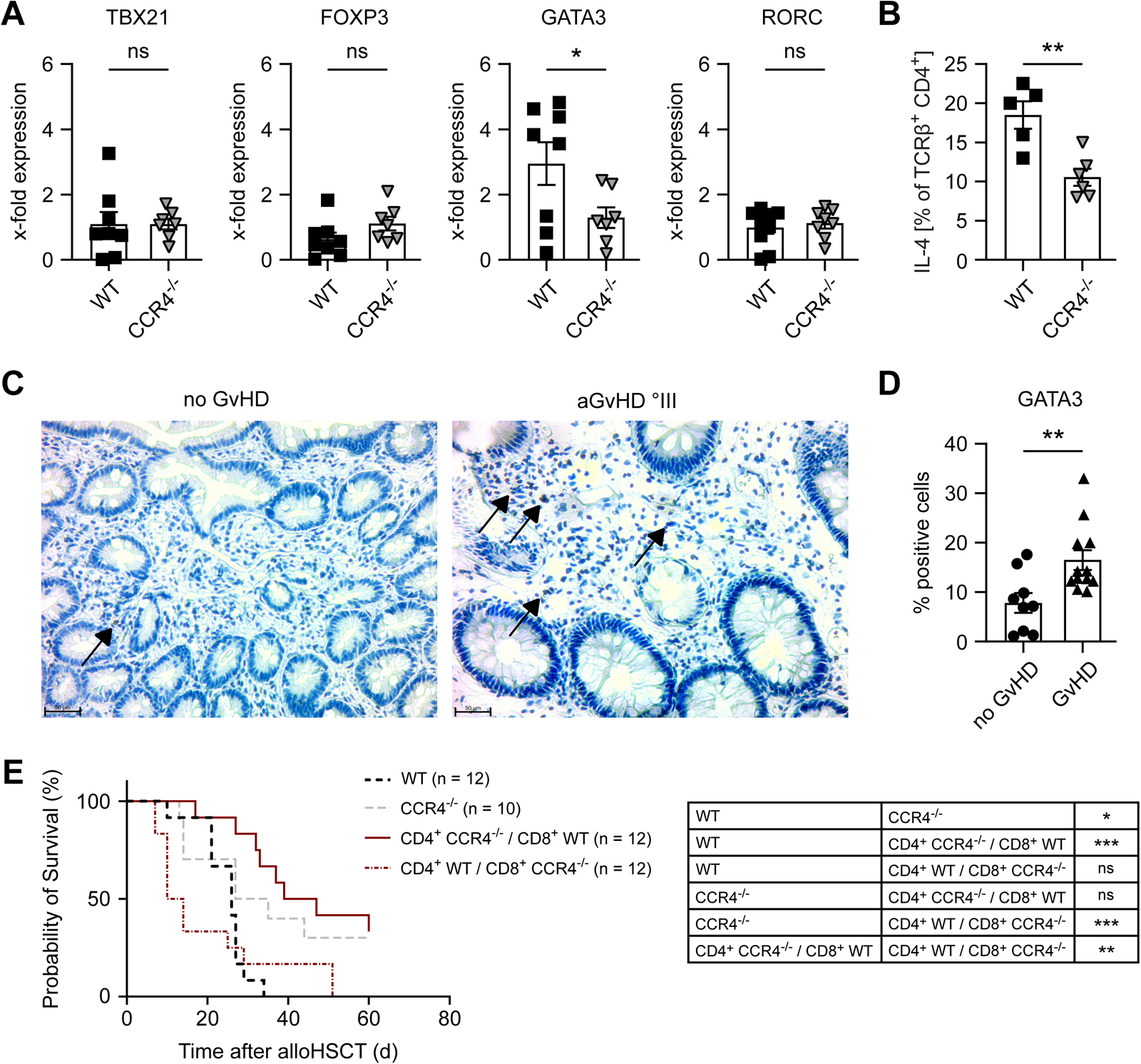
Transplantation of CCR4-/- reduces intestinal Th2 cells and lack of CCR4 only in the CD4 compartment mediates protection from GvHD. **(A)** Mice transplanted with WT or CCR4-/-T cells were sacrificed at day 7 after alloHSCT. RNA was isolated from whole small intestine. Levels of *Tbx21 (p=0.9894), Foxp3 (p=0.1228) Gata3 (p=0.0491) and Rorc (p=0.6236)* were quantified. (B) Leukocytes harvested seven days after alloHSCT were re-stimulated and IL-4 secretion was measured via FACS. Pooled statistical analysis from 2 independent experiments is shown; (*p=0.0031*). (C) Gut biopsies from patients without or with aGvHD °III were stained for GATA3. Arrows indicate Gata3-positive cells. (D) Percentage of Gata3-positive/all cells was calculated and decreased in patients withouth GvHD (no GvHD samples n=9, GvHD samples n=12; *p=0.0073*). (E) Mice received conditioning regimen and were transplanted with a physiological 3:2 mixture of CD4 and CD8 T cells, which were either proficient or deficient for CCR4. Mice transplanted with CD4 T cells lacking CCR4 showed prolonged survival. (CD4+ wt/ CD8+ wt vs. CD4+ CCR4-/-/ CD8+ CCR4-/- *p=0.0349*, CD4+ wt/ CD8+ wt vs. CD4+ CCR4-/-/ CD8+ wt *p=0.0001*, CD4+ wt/ CD8+ wt vs. CD4+ wt/ CD8+ CCR4-/- *p=0.5986*, CD4+ CCR4-/-/ CD8+ CCR4-/-vs. CD4+ CCR4-/-/ CD8+ wt *p=0.4909*, CD4+ CCR4-/-/ CD8+ CCR4-/-vs. CD4+ wt/ CD8+ CCR4-/- *p=0.0351*, CD4+ CCR4-/-/ CD8+ wt vs. CD4+ wt/ CD8+ CCR4-/-*p=0.0017*)

**Figure 3.**
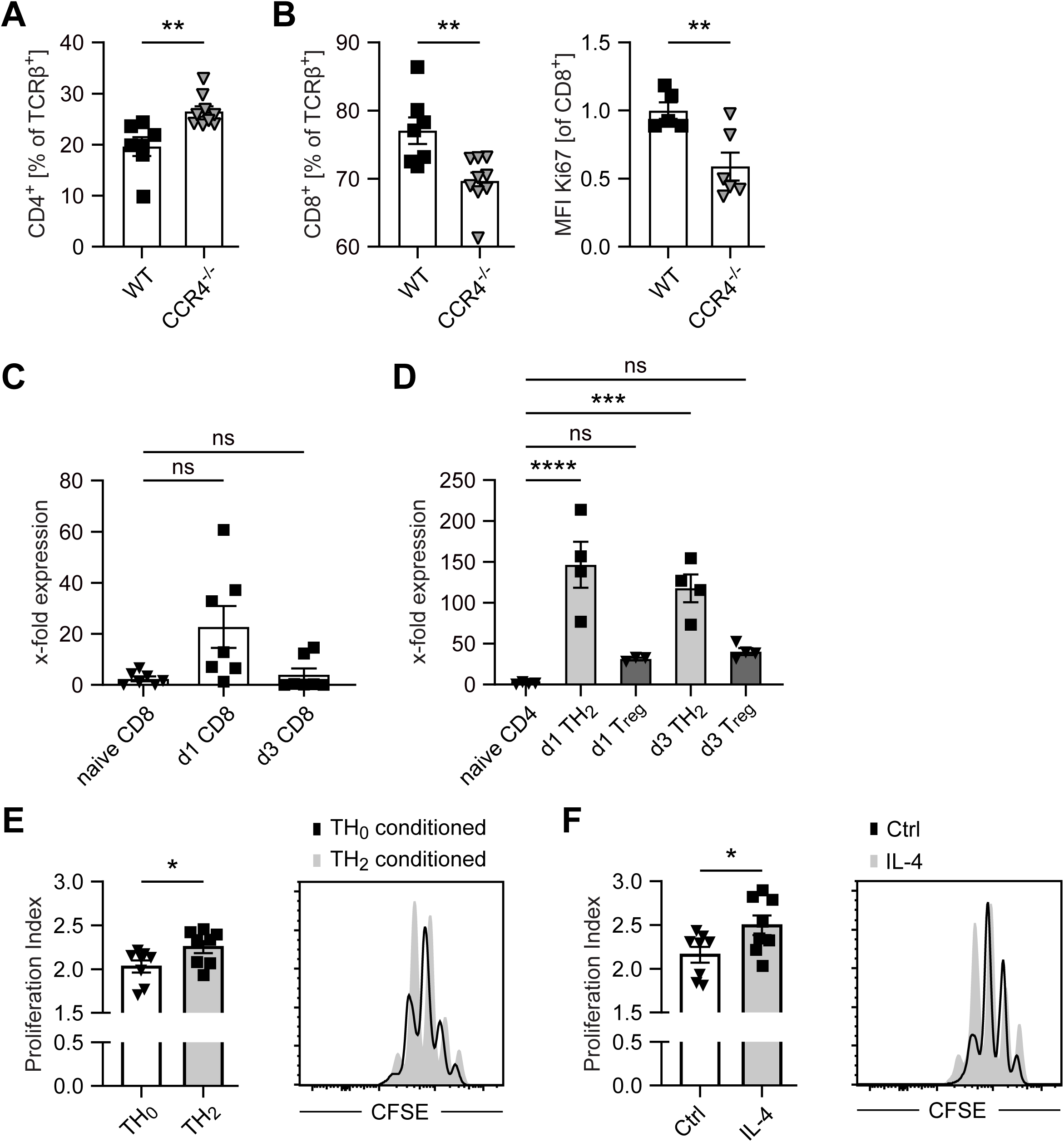
Only Th2 cells express relevant CCR4 and their cytokines foster CD8+ T cell proliferation. (A) On day 7 after alloHSCT leukocytes were isolated and stained for CD45, TCRβ, CD4 (*p=0.0038*) and (B) CD8 (*p=0.005*). Proliferation activity assessed by mean fluorescence intensity (MFI) of Ki67 was determined after permeabilization of isolated CD8+ T cells from the small intestine on day 7 after alloHSCT (*p=0.0097*). (C+D) Either naïve CD4+ or CD8+ were harvested directly after isolation or after 24h or 72h of proliferation in the presence of αCD3/CD28 beads. RNA was isolated and expression of CCR4 was determined via qPCR with the ΔΔCT method. Proliferating CD4+ T cells showed a significant upregulation of CCR4 after 24 h (*p<0.0001*) and 72 h (p=0.0004) in contrast to CD8+ T cells (*p=0.7723 after 24h and p>0.99 after 72h*). (E+F) Naïve splenic CD8+ cells were isolated and cultured in the presence of αCD3/CD28 beads. The proliferation of these cells, measured by their proliferation index, was enhanced in the presence of Th2 conditioned medium (E); (*p=0.0396*), or IL-4 (F);(*p=0.0344*).

These observations prompted us to analyze whether both CD4^+^ and CD8^+^ T cells upregulate CCR4 expression during proliferation. Therefore we measured the RNA expression of CCR4 in proliferating CD8^+^ T cells *in vitro*. We found that CD8^+^ T cells did not upregulate CCR4 24 or 72 h following aCD3/aCD28 stimulation (Fig. 3C, p=0.77 and p>0.99), suggesting that CD8^+^ T cells do not express relevant levels of CCR4.

We then assessed CCR4 expression in differentiating TH2 cells and Tregs. Compared to naïve CD4 T cells, TH2 cells exhibited significant transcriptional upregulation of CCR4 both 24 and 72 h after the start of differentiation.(Fig. 3D, p<0.0001 for 24h and p=0.0004 for 72h) Tregs also showed increased RNA levels of CCR4, albeit to a smaller, but not significant extent. These results support our data, indicating that CCR4^+^CD4^+^ T cells in aGvHD display a TH2 phenotype.

Based on our finding that intestinal CD8^+^ T cell proliferation is increased when CCR4^+^CD4^+^ T-cells are present (Fig. 3B), we assumed that TH2 cells might function as important triggers.

For this purpose, we performed *in vitro* CD8^+^ T cell proliferation in medium, which was either harvested from TH2-polarized or naïve CD4 T cell cultures, and found that the proliferation of CD8^+^ T cells was increased in TH2-conditioned medium (Fig. 3E, p=0.0396). Direct addition of IL-4 alone had similar effects (Fig. 3F, p=0.0344), corroborating our finding that elevated frequencies of intestinal IL-4^+^ CD4^+^ T cells correlated with increased frequencies of intestinal CD8^+^ T cells in T cell recipients.

### CD8^+^ T cell proliferation and migration is not CCR4-dependent, but CCL17 mediates CD4^+^ CCR4^+^ T cell migration *in vitro*

To further exclude the possibility that increased frequencies of CD8^+^ T cells are due to CCR4-dependent migration, we examined the effects of CCL17, a chemokine ligand for CCR4^25^, on CD8^+^ T cell migration and proliferation *in vitro*.

Using transwell assays, we demonstrated that CCL17 exclusively orchestrated the migration of CD4^+^ T cells in a CCR4-dependent manner, whereas the migration of CD8^+^ T cells was not CCL17-dependent (Fig. 4A, migration of wt CD3^+^ T cells (p=0.0046), wt CD4^+^ T cells (p=0.0035), and wt CD8^+^ T cells (p=0.3423) towards CCL17, Fig. 4B migration of CCR4^-/-^ CD3^+^ (p=0.1178), CD4^+^ (p=0.0662) and CD8^+^ (p=0.9742) T cells).

**Figure 4.**
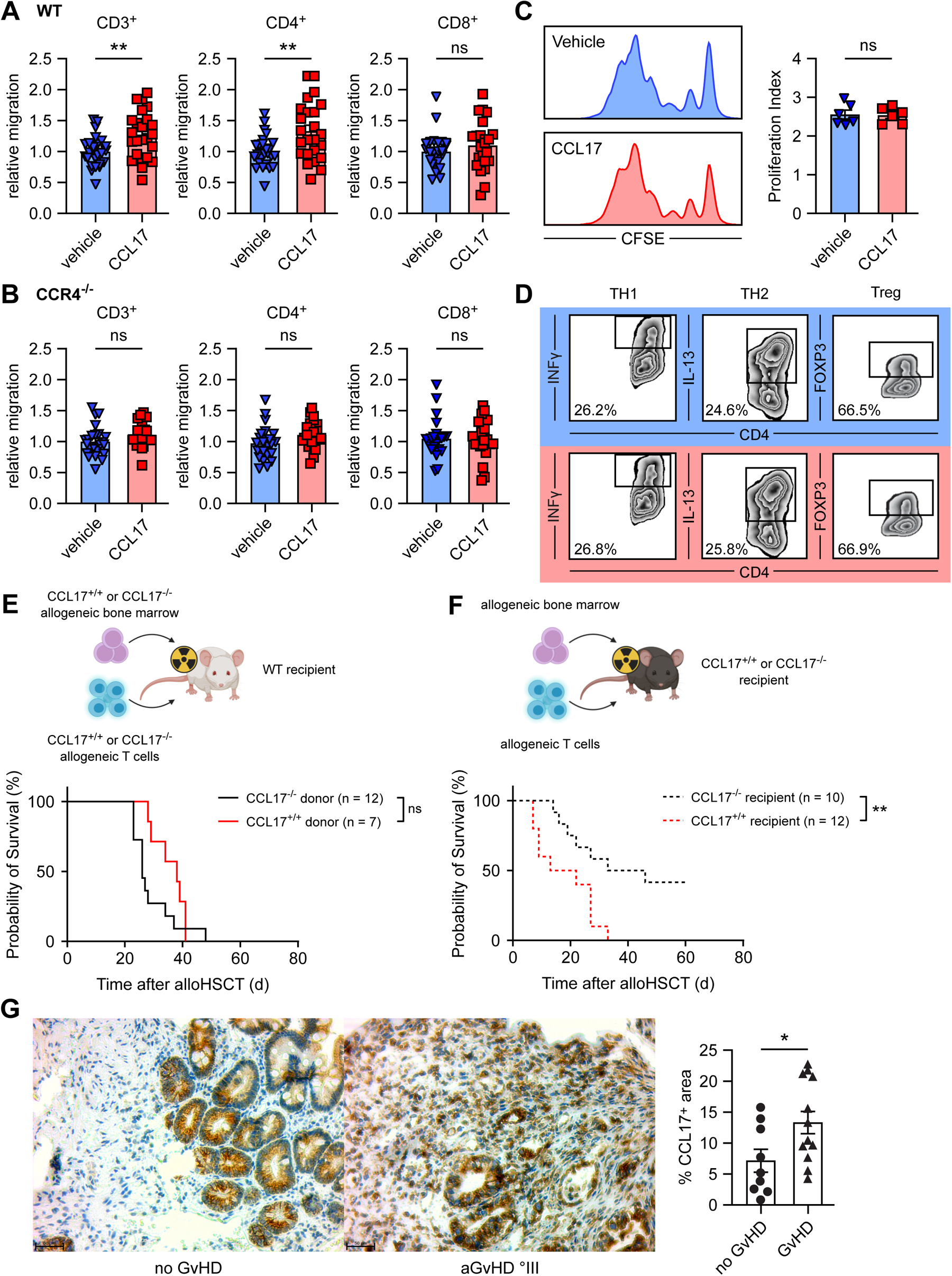
CCL17 only affects recruitment of CD4+ CCR4+ T cells and recipient CCL17 expression is associated with GvHD in mice and human. (A+B) Transwell migration assay of splenic T cells from either C57BL6/N wild-type mice or CCR4-/-animals was measured towards a CCL17 (800 ng/ml) gradient. Cells were counted via FACS for 90 seconds. WT CD3+ T cells (*p=0.0046*) and CD4+ T cells (*p=0.0035*) showed significant migration towards CCL17. (C) Naïve CD8+ T cells were isolated from the spleen of C57BL6/N mice. These cells were co-cultured with αCD3/CD28 beads and CCL17 (800ng/ml) when indicated. Proliferation was measured by CFSE distribution 60 hours later. (p *p=0.8986*) (C) Naïve CD4+ T cells were differentiated into Th1, Treg or Th2 cells in the presence (red) or absence of CCL17 (blue). Representative plots are shown. (D) Survival of BALB/c recipient mice after alloHSCT. Survival is not affected by CCL17 of donor bonemarrow or T cells; *p=0.11*. Schematic figure created using biorender.com. (E) CCL17-/-recipient mice showed prolonged survival compared to CCL17 +/+ recipients, *p=0.0069*. Schematic figure created using biorender.com. The data are pooled from 3 independent experiments with at least 3 mice per group. (F) Samples from gut biopsies of patients with the suspicion of intestinal aGvHD were collected. An experienced pathologist performed GvHD grading and samples were stained for CCL17 expression. Patients with proven aGvHD (n=12) show increased CCL17 positive area in representative areas compared to patients without GvHD (n=9). Representative samples of each group are shown.

Next, we stimulated naïve murine CD8^+^ T cells with αCD3/CD28 beads in the presence or absence of CCL17 and analyzed their proliferation; however, we detected no change (Fig. 4C, p=0.8986), indicating that CCL17 has no direct effects on CD8^+^ T cell proliferation.

We also assessed the effect of CCL17 on CD4^+^ Th differentiation *in vitro*. The frequencies of IFNγ^+^ (Th1), IL-13^+^ (TH2), and FOXP3^+^ (Treg) CD4^+^ T cells remained similar in the presence and absence of CCL17 (Fig. 4D).

These data suggest that CCL17 selectively orchestrates the migration of CD4^+^ CCR4^+^ T cells, specifically TH2 cells, which upregulate CCR4 upon proliferation (shown in Fig. 3D).

### CCL17 expression by the recipient and not the donor is relevant for aGvHD

Based on these results, we aimed to determine the functional role of CCL17 in experimental murine GvHD. We first transplanted recipient BALB/c mice with BM and T cells from either CCL17^+/+^ or CCL17^-/-^ C57BL/6 donors (Fig. 4E, top). CCL17 deficiency in the graft had no effect on overall survival, indicating that donor immune cell-derived CCL17 was not involved in aGvHD pathogenesis (Fig. 4E, p=0.11, Supplementary figure 1B for weight curves). In turn, when recipient CCL17^-/-^ mice received BM and T cells from WT donors (Balb/c → C57BL/6; Fig. 4F, top), they showed significantly reduced aGvHD-related mortality compared to wt recipients (Fig. 4F, p=0.0069 and Supplementary Figure 1E for weight curves), indicating that recipient-, and not donor-derived, CCL17 is involved in GvHD pathogenesis.

### CCL17 is upregulated in acute intestinal GvHD in humans

We subsequently analyzed the expression of CCL17 in our cohort of patients. Indeed, CCL17 was significantly upregulated in intestinal samples from patients who suffered from GvHD II°–IV°, in contrast to samples from patients without intestinal GvHD (Fig 4G, p=0.0307), corroborating our murine findings.

### CD11c^+^ cells are a relevant source for CCL17 in intestinal aGvHD *in vivo*

Analysis of CCL17 expression in our experimental mouse model revealed that RNA expression in the small intestine peaks approximately 3 days after alloHSCT (Supplementary Figure 1F, p=0.0038). DCs were shown to be crucially involved in aGvHD initiation^26^ and relevant sources of CCL17 are produced during intestinal inflammation in the context of Crohńs disease^27^. To investigate whether DCs are also a relevant source of CCL17 in the context of experimental murine aGvHD, we used CCL17 enhanced green fluorescent protein (eGFP) reporter mice (CCL17^eGFP/+^)^28^ as recipients (Balb/c → C57BL/6; Fig. 5A, top left). CCL17 eGFP reporter mice express enhanced green fluorescent protein (eGFP) under the control of the CCL17 promoter, which allows for visualization of CCL17-producing cell types. Cryosections of the small intestine of CCL17^eGFP/+^ recipients on day 3 post-transplantation were co-stained with CD11c, a prominent marker for identifying DCs.

**Figure 5.**
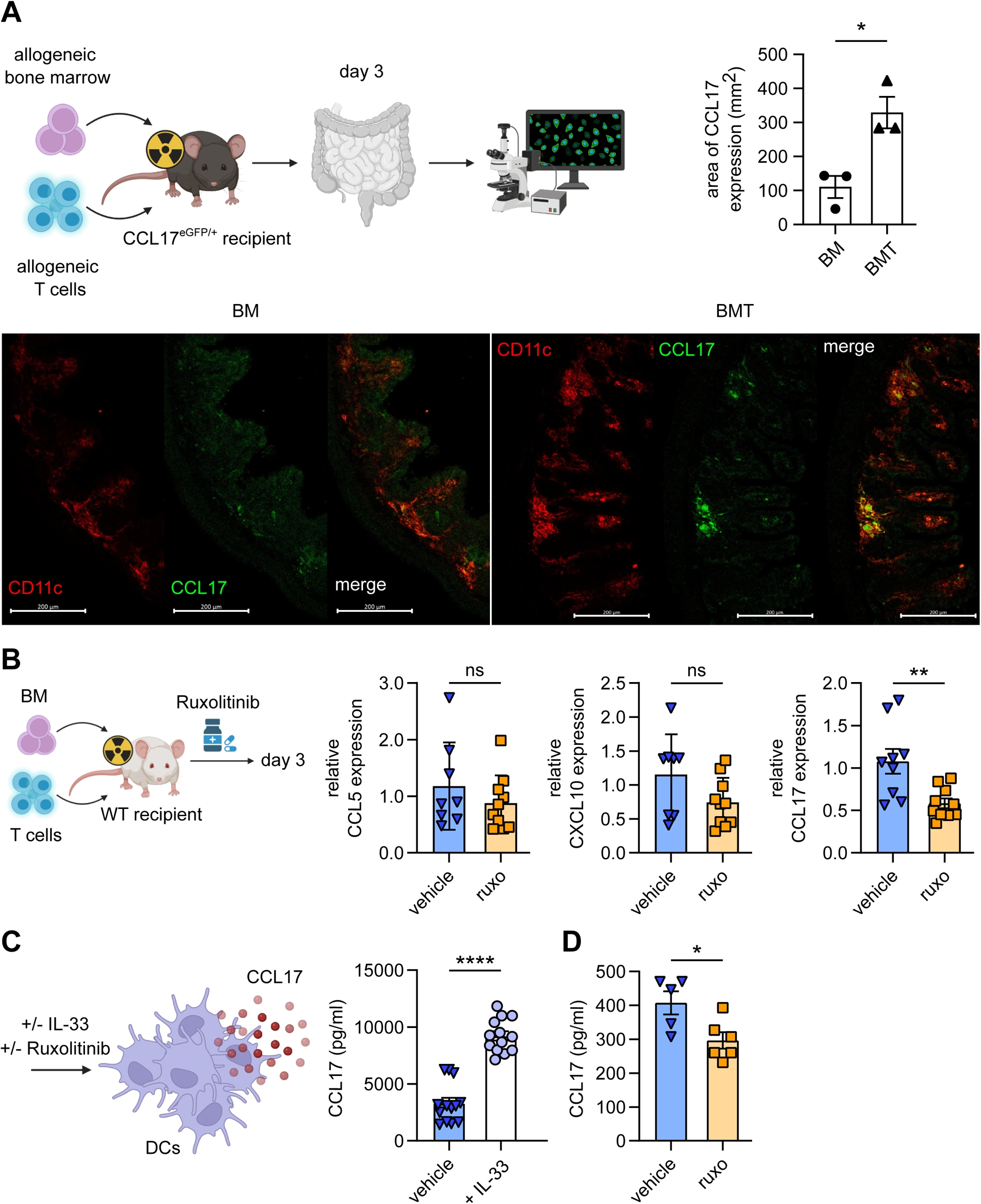
Host dendritic cells produce CCL17 after alloHSCT and ruxolitinib treatment reduces DC-derived CCL17 *in vitro* and intestinal CCL17 expression *in vivo*. (A) CCL17egfp/+ recipient mice underwent alloHSCT. Mice receiving only bone marrow cells (no aGvHD induction) were compared to those receiving additional T cells for induction of aGvHD. Quantification of the CCL17 positive area was higher in the GvHD group *p=0.027*. The experiment was performed twice. Representative data from one experiment is shown. Schematic figure created using biorender.com. Representative images of the small intestine stained for CD11c (DC marker) and DAPI are shown. Note the co-localization of CCL17(eGFP) and CD11c(PE). (B) Irradiated BALB/c mice received bone marrow cells and additional T cells from C57BL6/N donors. Mice were fed with ruxolitinib until day 3. Mice were sacrificed and CCL5 (*p=0.3267*), CXCL10 (*p=0.0622*) and CCL17 (*p=0.0045*) expression was determined via qPCR using the ΔΔCT method. Pooled data from 2 independent experiments. Schematic figure created using biorender.com. (C) Splenic DCs were treated with IL-33 for 18 hours, which induced CCL17 secretion (*p<0.0001).* (D) The presence of ruxolitinib (1µM) dampend CCL17 levels (*p=0.023*). Schematic figure created using

We found that CCL17 almost exclusively co-localized with CD11c (Fig 5A, bottom). In addition, the number of CD11c^+^ CCL17^+^ cells was markedly higher in aGvHD (BMT) mice than in non-GvHD (BM) mice (Fig. 5A top, p=0.027). Because only recipient-derived cells can express eGFP, these data provide evidence that intestinal host DCs are relevant sources of early intestinal CCL17 production *in vivo*.

### The JAK1/2 inhibitor ruxolitinib targets CCL17 expression in early intestinal aGvHD

Having shown that CCR4^+^ T cells might be harmful drivers of aGvHD, blocking CCR4 could be a promising strategy for preventing GvHD. However, CCR4 is also expressed on Tregs, which are known to be protective against aGvHD^1^., but CCR4-mediated antibody treatment may lead to severe colitis due to Treg depletion^29^, Therefore, targeting CCL17 may be a more successful and feasible approach for the treatment of GvHD.

In a previous study, we showed severe impairment of DC development, activation, migration, and cytokine production by the JAK1/2 inhibitor ruxolitinib^30^, without investigating its effects on CCL17 production. Since ruxolitinib has recently been approved for steroid-refractory acute and chronic GvHD^2,3,31^, we examined whether CCL17 could be affected by ruxolitinib treatment *in vivo*.

We adopted an established treatment protocol^20^ to our aGvHD model and administered ruxolitinib daily by oral gavage, starting one day prior to irradiation (Fig. 5B). We quantified the RNA expression of CCL17 and the T cell-associated chemokines CCL5 and CXCL10 in the intestine on day 3 after alloHSCT. CCL17 was significantly reduced in ruxolitinib-treated animals compared to that in vehicle-treated animals, while the effect of ruxolitinib on CXCL10 and CCL5 expression was less pronounced (Fig. 5C,CCL5 p=0.3267; CXCL10 p=0.0622; CCL17 p=0.0045).

Lastly, we confirmed that IL-33, a prominent alarmin released upon tissue damage, is a potent stimulus of CCL17 secretion in splenic DCs *in vitro*. Upon treatment with ruxolitinib, CCL17 release by DCs in response to IL-33 was significantly reduced (Fig. 5C, p=0.0230).

With these data, we provide additional insights into the immunosuppressive effects of ruxolitinib in GvHD and offer a therapeutic possibility to interfere with the CCR4-CCL17 axis in GvHD.

## Discussion

T cell-mediated tissue damage in target organs is a hallmark of aGvHD. Many studies have investigated various approaches to manipulate T-cell activation, proliferation, and function to prevent and treat GvHD. However, broad immunosuppression, diminished Graft-versus-Leukemia (GvL) activity, and increased relapse rates are common side effects that are detrimental to the success of alloHSCT. Interference with T cell migration has shown promising results in previous studies but has not yet been fully explored in the context of GvHD.

Here, we demonstrated that the CCR4-CCL17 axis may represent a novel target for early intervention in acute GvHD. We employed a standard MHC-mismatched mouse model for experimental aGvHD and investigated the roles of CCR4 and CCL17 by using genetic knockout mice. Our data showed that CCR4 expression, specifically on CD4^+^ donor T cells, significantly aggravated aGvHD severity and decreased overall survival in mice. Our findings correlate with the increased expression of GATA3 in the intestines of these mice as well as in human patients with aGvHD, accompanied by elevated IL-4 production by gut-infiltrating CD4^+^ T cells. Therefore, we hypothesized that CCR4^+^ CD4^+^ T cells that migrate into the intestine are predominantly TH2.

Direct tissue damage in GvHD target organs is mainly facilitated by CD8^+^ T cells, which express little to no CCR4, even after stimulation. Consequently, we found that the frequency of intestinal CD8^+^ T cells increased following transplantation of CCR4^+/+^ compared to CCR4^-/-^ T cells. These CD8^+^ T cells also displayed increased expression of the proliferation marker Ki67, and addition of either TH2-conditioned medium or IL-4 alone enhanced CD8^+^ T cell proliferation *in vitro*. Thus, our data suggest that the presence of CCR4^+^ CD4^+^ TH2 cells in the intestine fosters CD8^+^ T cell proliferation and effector functions, thereby promoting aGvHD pathology.

Our study supports previous evidence that both TH1 and TH2 cells are potent drivers of aGvHD (37). For example, IL-4 deficiency or the application of IL-4-blocking monoclonal antibodies protects against aGvHD ^32,33^. Elevated IL-13 serum levels were also associated with higher grade aGvHD ^34^ as seen in our setting, when reduced IL-13 serum levels in mice transplanted with CCR4^-/-^ T cells correlated with less severe aGvHD. In contrast, Helminth-induced IL-4 dampens aGvHD ^35^ and Zeiser et al. demonstrated that the administration of statins in experimental aGvHD induced a TH2 phenotype and reduced aGvHD lethality ^23^, suggesting a protective effect of TH2 cells. Based on these data and our own results, we speculate that TH2 polarization may play an ambivalent role in the context of aGvHD, possibly depending on the exact localization (e.g., lymphatic organ vs. end organ).

We also detected a decrease in serum levels of TH1-specific cytokines in mice that received CCR4^-/-^ T cells. These findings support the idea that both TH-subsets are hyper-activated in aGvHD and that CCR4-deficiency in donor T cells not only affects TH2-, but also dampens TH1-driven pathology. Similar observations were made in experimental TH1-dependent models of inflammatory bowel disease (IBD), in which IL-4 exacerbated the disease severity ^36^.

The existing prophylactic and therapeutic strategies for aGvHD are largely based on immunosuppression by modulating T-cell proliferation ^1^. Disruption of chemokine receptor-mediated T cell function has also shown promising results ^11^, but interference with the CCR4 receptor, in particular, has not been investigated in aGvHD thus far. Vorinostat, a histone deacetylase (HDAC) inhibitor, was found to be effective in preventing aGvHD ^37^ by modulating DC cytokine production and balancing circulating T cell subsets towards a more anti-inflammatory setting (TH1/TH17 ↓, Tregs↑)^38^. Interestingly, it has been shown that it also downregulates CCR4 expression on T cells ^39^, which may mechanistically explain its mode of action and is in line with our data showing GvHD improvement when CCR4 is absent in T cells.

Importantly, interference with the CCR4 receptor must be carefully examined. In ulcerative colitis, adoptively transferred CCR4-deficient Tregs fail to inhibit T cell proliferation due to delayed migration ^40^. In turn, treatment of adult T-cell lymphoma with the CCR4 antibody mogamulizumab can cause colitis mimicking GvHD, which is suspected to be due to Treg depletion ^29^. Therefore, the protective capacity of Tregs should be considered when targeting CCR4 in GvHD.

Instead, modulating chemokine, and not chemokine receptor, expression may be a more elaborate approach. CCL17, a potent ligand of CCR4, has previously been implicated in intestinal inflammation. For example, in IBD, CCL17 counteracted Treg-mediated protection ^13^ and CCL17 deficiency enhanced allograft survival after solid organ transplantation ^28^. We show that CCL17 expression is upregulated in biopsies of patients with acute intestinal GvHD, and CCL17 expression in recipient but not in donor cells correlated with increased aGvHD lethality in our mouse model. We further identified recipient-derived CD11c^+^ DCs as the most relevant source of CCL17 in the intestine of mice following alloHSCT. Interestingly, pharmacological interventions known to improve experimental GvHD, such as statins ^23^, repress CCL17 expression by DCs ^41^ and the protective effect of statins is less pronounced in TH2-independent models. Therefore, protection from GvHD by statins may be at least partially explained by the modulation of CCL17-mediated TH2 cell trafficking to the GI tract ^23^.

Furthermore, we showed that IL-33, one of the early alarmins released upon conditioning-induced tissue damage, could stimulate CCL17 production in DCs *in vitro*. Although the ambivalent role of IL-33 has been extensively discussed in the context of T cell activation *via* the suppression of tumorigenicity 2 (ST2) in aGvHD ^42,43^, its direct effects on DC biology in aGvHD have not been defined. Some data also suggest that IL-33 can act as a chemoattractant for TH2 cells^44^.

Recently, the JAK1/2 inhibitor, ruxolitinib, was approved for the treatment of steroid-refractory GvHD. We have previously shown that ruxolitinib suppresses DC activation and function^20,45^ and its pleiotropic immunosuppressive effects have been reported in multiple studies ^30,30,45–48^. Here, we show that ruxolitinib also inhibits IL-33-induced CCL17 secretion by DCs *in vitro* and reduces CCL17 expression in the intestines of mice immediately after alloHSCT. These results may not only add to the understanding of how ruxolitinib specifically combats GvHD, but also provide a basis for future designs of targeted therapies.

## Conclusion

We propose that the early release of IL-33 upon conditioning-induced tissue damage in the GI tract stimulates CCL17 secretion by host-derived dendritic cells. CCR4^+^ donor T cells, predominantly of the TH2 phenotype, subsequently infiltrate the intestines as a GvHD target organ and promote the expansion of CD8^+^ T cells. The initial upregulation of CCL17 secretion may be suppressed by the JAK1/2 inhibitor, ruxolitinib, thereby indirectly disrupting the CCR4-CCL17 axis. Overall, our data shed light on the role of CCR4 and CCL17 in acute GvHD and present versatile opportunities for targeting chemotaxis as a prophylactic and therapeutic treatment strategy.

## Ethics approval and consent to participate

This study was conducted in accordance with the Declaration of Helsinki, and approval was obtained from the Institutional Ethics Committee of the University of Bonn (#175/20). All animal experiments were approved by *Landesamt für Natur, Umwelt und Verbraucherschutz Nordrhein-Westfalen (LANUV)*.

## Consent for publication

The authors declare no competing interests.

## Availability of data and materials

The datasets used and/or analyzed during the current study are available from the corresponding author upon reasonable request.

## Authorśdisclousure

The authors declare that they have no competing interests.

## Authors’ contributions

S.S., S.K.J., M.K., J.B-G., C.F., and O.S. performed the experiments, analyzed the results, and made the figures; G.K. and M.T. helped to investigate human biopsies. S.S., A.H., M.K., and D.W. designed the research and wrote the paper; P.B. and C.K. designed the research and discussed the results.

## Acknowledgements

The authors thank Anja Schmidt, Vadim Kotov, and Susanne Steiner for technical assistance. This work was supported by BONFOR-Forschungsförderung (O-142.0016 and O-142-0019 to S.S., and O-142.0024 to O.S.). Additionally, this work was funded by Deutsche Krebshilfe through a Mildred Scheel Nachwuchszentrum Grant (Grant number 70113307 to S.S.) and supported by a grant from Deutsche Forschungsgemeinschaft (DFG EXC2151–390873048 to A.H.).

---

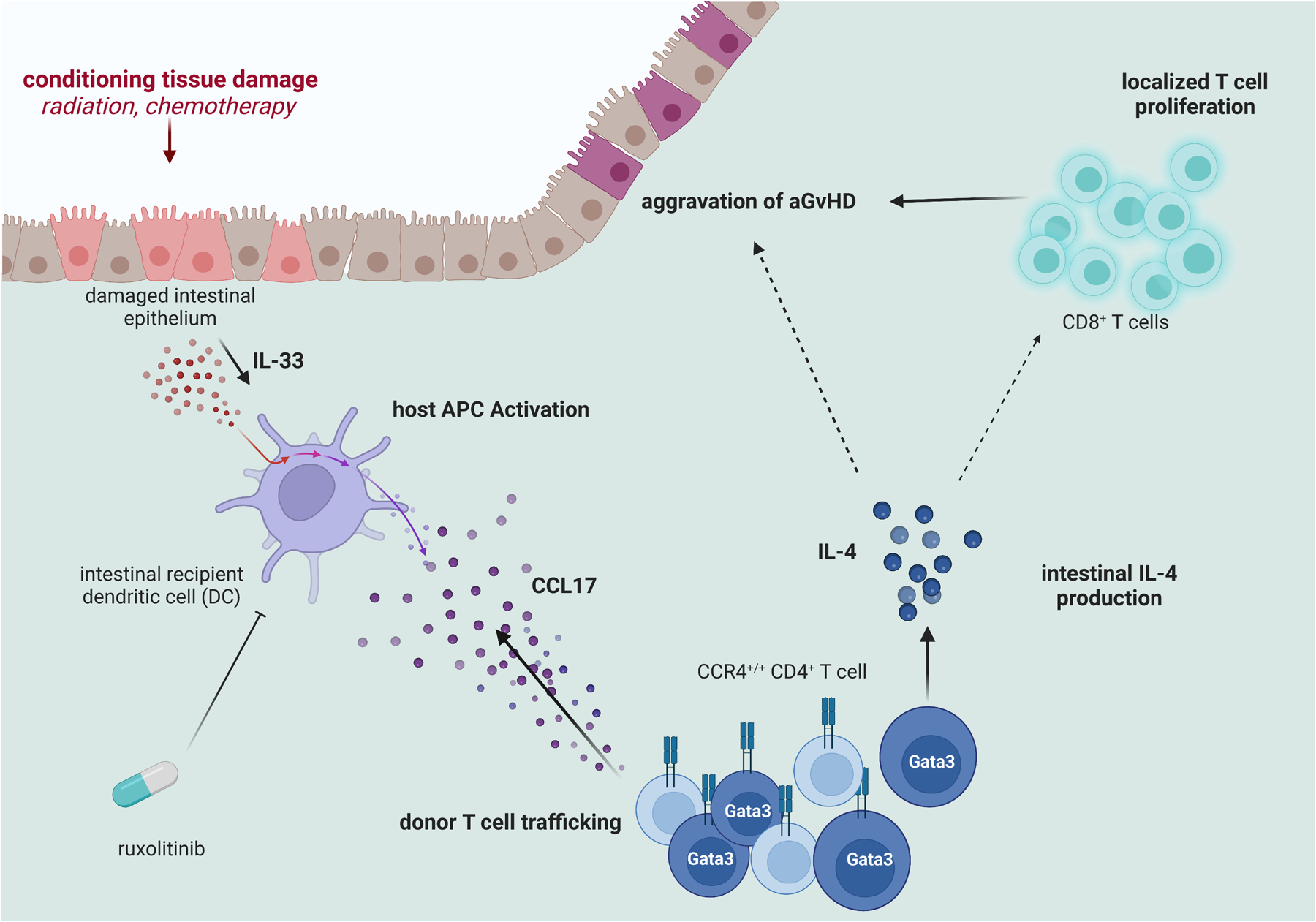

## Notes

### Competing Interest Statement

The authors have declared no competing interest.

## References

1. Zeiser, R. & Blazar, B. R. Acute Graft-versus-Host Disease — Biologic Process, Prevention, and Therapy. N. Engl. J. Med. 377, 2167–2179 (2017).

2. Zeiser, R. et al. Ruxolitinib for Glucocorticoid-Refractory Acute Graft-versus-Host Disease. N. Engl. J. Med. 382, 1800–1810 (2020).

3. Zeiser, R. et al. Ruxolitinib for Glucocorticoid-Refractory Chronic Graft-versus-Host Disease. N. Engl. J. Med. 385, 228–238 (2021).

4. Schwab, L. et al. Neutrophil granulocytes recruited upon translocation of intestinal bacteria enhance graft-versus-host disease via tissue damage. Nat. Med. 20, 648–654 (2014).

5. Zeiser, R., Socié, G. & Blazar, B. R. Pathogenesis of acute graft-versus-host disease: from intestinal microbiota alterations to donor T cell activation. Br. J. Haematol. 175, 191–207 (2016).

6. Zlotnik, A. & Yoshie, O. The Chemokine Superfamily Revisited. Immunity 36, 705–716 (2012).

7. Carbone, F. R., Kurts, C., Bennett, S. R. M., Miller, J. F. A. P. & Heath, W. R. Cross-presentation: a general mechanism for CTL immunity and tolerance. Immunol. Today 19, 368–373 (1998).

8. Wysocki, C. A., Panoskaltsis-Mortari, A., Blazar, B. R. & Serody, J. S. Leukocyte migration and graft-versus-host disease. Blood 105, 4191–4199 (2005).

9. Wysocki, C. A. et al. Differential Roles for CCR5 Expression on Donor T Cells during Graft-versus-Host Disease Based on Pretransplant Conditioning. J. Immunol. 173, 845–854 (2004).

10. Piper, K. P. et al. CXCL10-CXCR3 interactions play an important role in the pathogenesis of acute graft-versus-host disease in the skin following allogeneic stem-cell transplantation. Blood 110, 3827–3832 (2007).

11. Moy, R. H. et al. Clinical and immunologic impact of CCR5 blockade in graft-versus-host disease prophylaxis. Blood 129, 906–916 (2017).

12. Ait Yahia, S., et al. CCL17 production by dendritic cells is required for NOD1-mediated exacerbation of allergic asthma. Am. J. Respir. Crit. Care Med. 189, 899–908 (2014).

13. Heiseke, A. F. et al. CCL17 Promotes Intestinal Inflammation in Mice and Counteracts Regulatory T Cell–Mediated Protection From Colitis. Gastroenterology 142, 335–345 (2012).

14. Ruland, C. et al. Chemokine CCL17 is expressed by dendritic cells in the CNS during experimental autoimmune encephalomyelitis and promotes pathogenesis of disease. Brain. Behav. Immun. 66, 382–393 (2017).

15. Zhou, L., Askew, D., Wu, C. & Gilliam, A. C. Cutaneous Gene Expression by DNA Microarray in Murine Sclerodermatous Graft-Versus-Host Disease, a Model for Human Scleroderma. J. Invest. Dermatol. 127, 281–292 (2007).

16. Imai, T. et al. The T Cell-directed CC Chemokine TARC Is a Highly Specific Biological Ligand for CC Chemokine Receptor 4. J. Biol. Chem. 272, 15036–15042 (1997).

17. Imai, T. et al. Macrophage-derived Chemokine Is a Functional Ligand for the CC Chemokine Receptor 4. J. Biol. Chem. 273, 1764–1768 (1998).

18. Kim, S. H., Cleary, M. M., Fox, H. S., Chantry, D. & Sarvetnick, N. CCR4-bearing T cells participate in autoimmune diabetes. J. Clin. Invest. 110, 1675–1686 (2002).

19. Palchevskiy, V. et al. CCR4 expression on host T cells is a driver for alloreactive responses and lung rejection. JCI Insight 5, e121782, 121782 (2019).

20. Heine, A. et al. The JAK-inhibitor ruxolitinib impairs dendritic cell function in vitro and in vivo. Blood 122, 1192–1202 (2013).

21. Hülsdünker, J. & Zeiser, R. In Vivo Myeloperoxidase Imaging and Flow Cytometry Analysis of Intestinal Myeloid Cells. Methods Mol. Biol. Clifton NJ 1422, 161–167 (2016).

22. Reshef, R. et al. Extended CCR5 Blockade for Graft-versus-Host Disease Prophylaxis Improves Outcomes of Reduced-Intensity Unrelated Donor Hematopoietic Cell Transplantation: A Phase II Clinical Trial. Biol. Blood Marrow Transplant. J. Am. Soc. Blood Marrow Transplant. 25, 515–521 (2019).

23. Zeiser, R. et al. Preemptive HMG-CoA reductase inhibition provides graft-versus-host disease protection by Th-2 polarization while sparing graft-versus-leukemia activity. Blood 110, 4588– 4598 (2007).

24. Imai, T. et al. Selective recruitment of CCR4-bearing Th2 cells toward antigen-presenting cells by the CC chemokines thymus and activation-regulated chemokine and macrophage-derived chemokine. Int. Immunol. 11, 81–88 (1999).

25. Semmling, V. et al. Alternative cross-priming through CCL17-CCR4-mediated attraction of CTLs toward NKT cell–licensed DCs. Nat. Immunol. 11, 313–320 (2010).

26. Chen, S. et al. MicroRNA-155-deficient dendritic cells cause less severe GVHD through reduced migration and defective inflammasome activation. Blood 126, 103–112 (2015).

27. Martin, J. C. et al. Single-Cell Analysis of Crohn’s Disease Lesions Identifies a Pathogenic Cellular Module Associated with Resistance to Anti-TNF Therapy. Cell 178, 1493–1508.e20 (2019).

28. Alferink, J. et al. Compartmentalized Production of CCL17 In Vivo. J. Exp. Med. 197, 585–599 (2003).

29. Ishitsuka, K. et al. Colitis mimicking graft-versus-host disease during treatment with the anti-CCR4 monoclonal antibody, mogamulizumab. Int. J. Hematol. 102, 493–497 (2015).

30. Heine, A., Brossart, P. & Wolf, D. Ruxolitinib is a potent immunosuppressive compound: is it time for anti-infective prophylaxis? Blood 122, 3843–3844 (2013).

31. Jagasia, M. et al. Ruxolitinib for the treatment of steroid-refractory acute GVHD (REACH1): a multicenter, open-label phase 2 trial. Blood 135, 1739–1749 (2020).

32. Murphy, W. J. et al. Differential effects of the absence of interferon-gamma and IL-4 in acute graft-versus-host disease after allogeneic bone marrow transplantation in mice. J. Clin. Invest. 102, 1742–1748 (1998).

33. Ushiyama, C. et al. Anti-IL-4 antibody prevents graft-versus-host disease in mice after bone marrow transplantation. The IgE allotype is an important marker of graft-versus-host disease. J. Immunol. 154, 2687–2696 (1995).

34. Jordan, W. J. IL-13 production by donor T cells is prognostic of acute graft-versus-host disease following unrelated donor stem cell transplantation. Blood 103, 717–724 (2004).

35. Li, Y. et al. Helminth-Induced Production of TGF-β and Suppression of Graft-versus-Host Disease Is Dependent on IL-4 Production by Host Cells. J. Immunol. Baltim. Md 1950 201, 2910–2922 (2018).

36. Bamias, G. et al. Proinflammatory effects of TH2 cytokines in a murine model of chronic small intestinal inflammation. Gastroenterology 128, 654–666 (2005).

37. Choi, S. W. et al. Vorinostat plus tacrolimus/methotrexate to prevent GVHD after myeloablative conditioning, unrelated donor HCT. Blood 130, 1760–1767 (2017).

38. Holtan, S. G. & Weisdorf, D. J. Vorinostat is victorious in GVHD prevention. Blood 130, 1690–1691 (2017).

39. Kitadate, A. et al. Histone deacetylase inhibitors downregulate CCR4 expression and decrease mogamulizumab efficacy in CCR4-positive mature T-cell lymphomas. in Haematologica (2018). doi:10.3324/haematol.2017.177279.

40. Yuan, Q. et al. CCR4-dependent regulatory T cell function in inflammatory bowel disease. J. Exp. Med. 204, 1327–1334 (2007).

41. Inagaki-Katashiba, N. et al. Statins can suppress DC-mediated Th2 responses through the repression of OX40-ligand and CCL17 expression. Eur. J. Immunol. 49, 2051–2062 (2019).

42. Reichenbach, D. K. et al. The IL-33/ST2 axis augments effector T-cell responses during acute GVHD. Blood 125, 3183–3192 (2015).

43. Matta, B. M. et al. Peri-alloHCT IL-33 administration expands recipient T-regulatory cells that protect mice against acute GVHD. Blood 128, 427–439 (2016).

44. Komai-Koma, M. et al. IL-33 is a chemoattractant for human Th2 cells. Eur. J. Immunol. 37, 2779– 2786 (2007).

45. Rudolph, J. et al. The JAK inhibitor ruxolitinib impairs dendritic cell migration via off-target inhibition of ROCK. Leukemia 30, 2119–2123 (2016).

46. Parampalli Yajnanarayana, S., et al. JAK1/2 inhibition impairs T cell function in vitro and in patients with myeloproliferative neoplasms. Br. J. Haematol. 169, 824–833 (2015).

47. Schönberg, K. et al. JAK Inhibition Impairs NK Cell Function in Myeloproliferative Neoplasms. Cancer Res. 75, 2187–2199 (2015).

48. Elli, E. M., Baratè, C., Mendicino, F., Palandri, F. & Palumbo, G. A. Mechanisms Underlying the Anti-inflammatory and Immunosuppressive Activity of Ruxolitinib. Front. Oncol. 9, 1186 (2019).

