## Supplementary Material for "The CCR4/CCL17 axis drives intestinal acute Graft versus Host disease after allogeneic bone marrow transplantation"

**Supplementary Table 1: Patient characteristics**

| # | GvHD ° | reason for transplant | day post alloHSCT | age | conditioning regime | immunosuppression |
|---|---|---|---|---|---|---|
| 1 | no | t-MDS | 20 | 21 | TT/Bu/Flu | pt-Cy/ Tac/ MMF |
| 2 | no | CNL | 216 | 65 | Flu/Amsa/ARA-C/Treo/Cy/ATG | CsA/MMF |
| 3 | no | sAML | 28 | 70 | Flu/BCNU/Mel/ATG | CsA/MMF |
| 4 | no | sAML | 108 | 71 | Flu/BCNU/Mel | CsA/MMF |
| 5 | no | AML | 19 | 59 | Flu/BCNU/Mel/ATG | Tac/MMF |
| 6 | no | MDS/MPN | 141 | 45 | Bu/Cy/ATG | Tac/MMF |
| 7 | no | MDS | 105 | 72 | Flu/Bu | Tac/MMF/Sir |
| 8 | no | AML | 177 | 63 | Flu/BCNU/Mel | CsA/MMF |
| 9 | no | MDS/MPN | 217 | 41 | TT/Bu/Flu | pt-Cy/ Tac/ MMF/ CsA |
| | | | 114,5555556 | | | |
| 11 | 4 | Mantle cell lymphoma | 35 | 58 | Treo/Flu | CsA/MMF |
| 12 | 3 | sAML | 98 | 56 | Flu/Cy/TBI | pt-Cy/Tac/MMF |
| 13 | 4 | sAML | 68 | 58 | Flu/BCNU/Mel | CsA/MMF |
| 14 | 4 | MDS | 123 | 54 | Bu/Flu | CsA/MMF |
| 15 | 4 | AML | 32 | 72 | Flu/BCNU/Mel | CsA/MMF |
| 16 | 3 | AML | 59 | 54 | Flu/BCNU/Mel | CsA/MMF |
| 17 | 3 | sAML | 54 | 57 | Mel/Flu/Cy/TBI | pt-Cy/Tac/MMF/ CsA |
| 18 | 3 | OMF | 222 | 29 | Flu/TBI/ATG | CsA/MTX |
| 19 | 4 | MDS | 21 | 39 | Flu/Treo/ATG | CsA/MMF/Tac |
| 20 | 2 | AML | 162 | 59 | Flu/BCNU/Mel | CsA/MMF |
| 21 | 4 | AML | 57 | 45 | Mel/Flu/Tre | CsA/MMF |
| 22 | 3 | MDS | 172 | 72 | Flu/Cy/Mel/TBI | pt-Cy/Tac/MMF |
| 23 | 3 | AML | 138 | 62 | TT/Bu/Flu | pt-Cy/Tac.MMF |
| | | | 95,46153846 | | | |

Supplementary Fig. 1

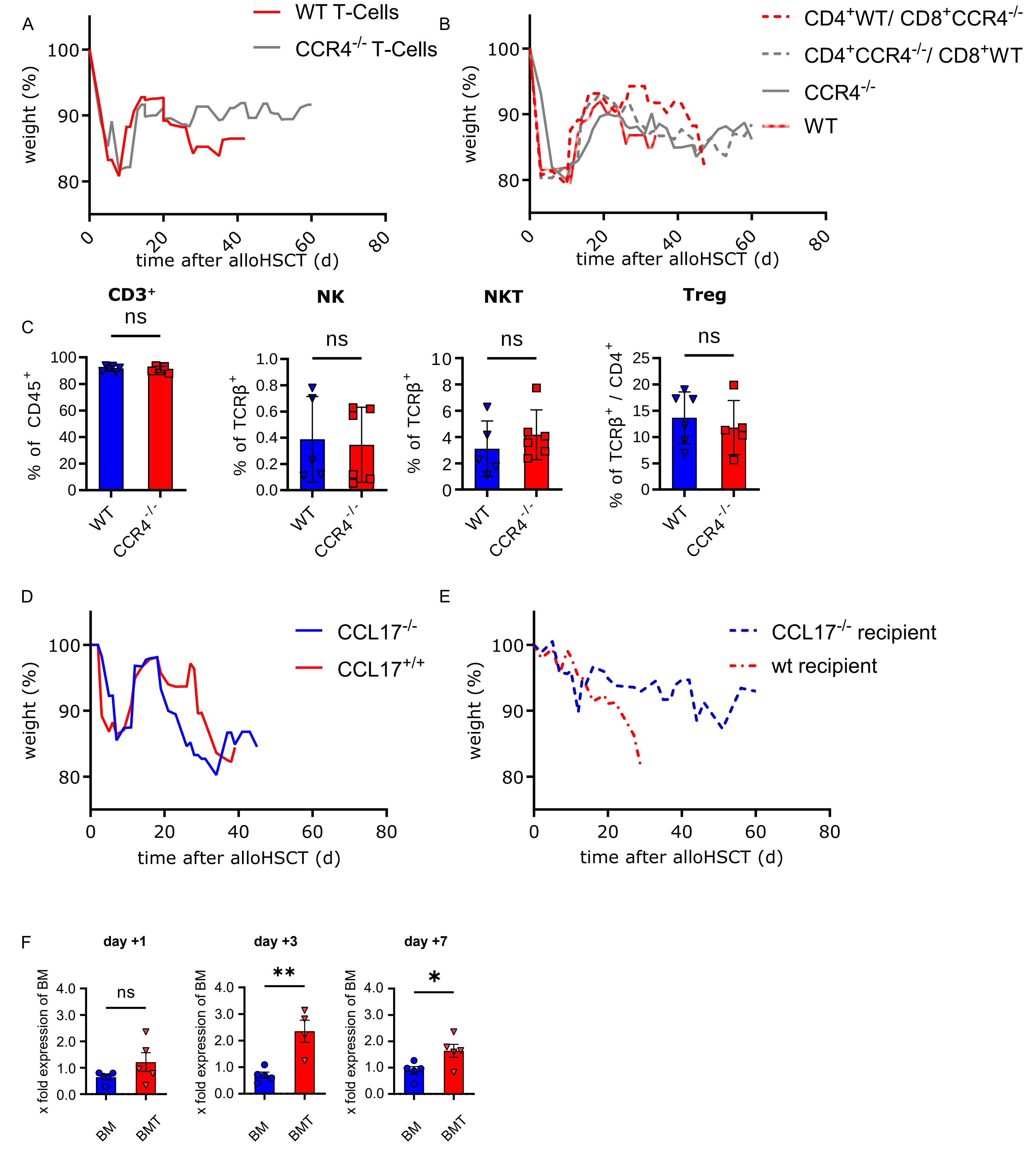

### Supplementary Figure legends

(A) Balb/c mice were transplanted with bone marrow from wt mice and T cell proficient (wt) or deficient for CCR4 (CCR4$^{-/-}$). Corresponding to the Kaplan-Meier plot in Fig 1A, weight was measured and depicted. Recipients of wt T cells show increased weight loss as a sign of aGvHD.

(B) Balb/c mice were transplanted with CD4$^{+}$ and CD8$^{+}$ T cells from either CCR4$^{-/-}$ or wt donor mice in a physiological 3:2 ratio. Only mice lacking CCR4 in the CD4$^{+}$ compartment (CCR4$^{-/-}$ and CD4$^{+}$ CCR4$^{-/-}$/CD8$^{+}$ wt, red lines) show recovery after initial weight loss until the end of the experiment. As an additional proof of clinical GvHD mice receiving wt CD4$^{+}$ T cells (wt and CD4$^{+}$ wt/CD8$^{+}$ CCR4$^{-/-}$, grey lines) show pronounced weight loss.

(C) Balb/c recipient mice, transplanted with either wt (blue) or CCR4$^{-/-}$ (red) T cells, were sacrificed 7 days after transplantation. Small intestines were isolated and infiltrating T cells were analyzed by FACS. CD45$^{+}$ allogeneic T cells were analyzed and CD3 (p=0.6605), NK (p=0.8277), NKT (p=0.3426) and Treg (p=0.1059) cells did not differ.

(D) Recipients of CCL17$^{-/-}$ and wt T cells and BM show comparable loss of weight after induction of murine experimental aGvHD.

(E) CCL17$^{-/-}$ recipients present with less severe loss of weight after irradiation and transplantation of T cells compared to wt recipients.

(F) Balb/c recipient mice were irradiated and transplanted with bone marrow cells from C57BL6/N donor mice (BM). T cells were additionally transplanted in the experimental group (BMT). Mice were sacrificed after 1, 3 and 7 days. RNA from the terminal ileum was isolated. CCL17 expression was calculated by the ΔΔCT method using GAPDH as a housekeeping gene. Pooled data from 2 experiments. Expression on day 3 and on day 7 was significantly increased; *p=0.0038 and p=0.03*.

### Supplementary Methods

#### Time depedent analysis of CCL17 expression assay

Balb/c mice were irradiated and received bone marrow cells from wt mice (BM group). Additional T cells were injected to induce aGvHD (BMT). Mice were sacrificied after 1,3 and 7 days and the small intestine was removed and snap frozen in liquid nitrogen. RNA was isolated by RNAzol (Sigma Aldrich, #R4533). Using the RevertAid First Strand cDNA Synthesis Kit (Thermofisher, #18091050) according to the manufacturer's instructions, cDNA was synthesized and quantitative real-time PCR (qPCR) was performed using specific primers at 0.2µM concentration and SYBR Green PCR Master Mix (Applied Biosystems, #4309155) on the Mastercycler RealPlex2 (Eppendorf). Data were normalized to GAPDH expression and analyzed using the ΔΔCT method. Primer sequence is listed in supplementary table 2.

Supplementary table 2

Primer sequences

| | |
|---|---|
| CCL5 | 5′-TGCTGCTTTGCCTACCTCTC-3′ 5′-TCCTTCGAGTGACAAACA CGA-3′ |
| CXCL10 | 5′-TCATCCTGCTGGGTCTGAGT-3′; 5′-ATCGTGGCAATGATCTCAACAC-3′ |
| CCL17 | 5′-TGGTATAAGACCTCAGTGGAGTGTTC-3′, 5′-GCTTGCCCTGGACAGTCAGA-3′ |
| CCR4 | 5´-CTTTCAGAAGAGCAAGGCAGCTC-3´ 5´-ATCTTTGGAATCGTCGCGCT-3´ |

FACS antibodies

| | |
|---|---|
| CD4 | clone GK1.5 |
| CD8a | clone 53-6.7 |
| CD11b | clone M1/70 |
| FOXP3 | clone FJK-16s |
| H-2Dd | clone 34-2-12 |
| IL-4 | clone 11B11 |
| KI67 | clone 16A8 |
| Live/Dead | Invitrogen, #L34965 |
| NK1.1, | clone PK136 |
| TCRß | clone H57-597 |

Th1 medium

| | |
|---|---|
| IL12 | 5 ng/ml, Peprotech, #210-12 |
| IL2 | 50 units/ml, Peprotech, #200-02 |
| anti-IL4 | 10 µg/ml, Biolegend, #504122 |

Th2 medium

| | |
|---|---|
| IL-4, | 10 ng/ml, Peprotech, #214-14 |
| IL-2 | 10 ng/ml, Peprotech, #210-12 |
| anti-IL12 | 10 µg/ml, Biolegend, #505308 |

iTreg medium

| | |
|---|---|
| TGF-β | 5 ng/ml, Peprotech, #100-21 |
| IL2 | 100 units/ml, Peprotech, #200-02 |

stimulation medium

| | |
|---|---|
| PMA | 50ng/ml, Sigma-Aldrich, #P8139 |
| Ionomycin | 1µg/ml, Sigma Aldrich, # I9657 |
| Brefeldin A | 1000x, Biolgend, #420601 |
| Monensin | 1000x, Biolegend, #420701 |
